# Seasonal variation of genotypes and reproductive plasticity in a facultative clonal freshwater invertebrate animal (Hydra oligactis) living in a temperate lake

**DOI:** 10.1101/2021.09.16.460593

**Authors:** Máté Miklós, Levente Laczkó, Gábor Sramkó, Zoltán Barta, Jácint Tökölyi

## Abstract

Facultative sexual organisms combine sexual and asexual reproduction within a single life cycle, often switching between reproductive modes depending on environmental conditions. These organisms frequently inhabit variable seasonal environments, where favourable periods alternate with unfavourable periods, generating temporally varying selection pressures that strongly influence life history decisions and hence population dynamics. Due to the rapidly accelerating changes in our global environment today, understanding the dynamics of and genetic changes in facultative sexual populations inhabiting seasonal environments is critical to assess and prepare for additional challenges that will affect such ecosystems. In this study we aimed at obtaining insights of the seasonal population dynamics of the facultative sexual freshwater cnidarian *Hydra oligactis* through a combination of Restriction-site Associated Sequencing (RAD-Seq) genotyping and the collection of phenotypic data on the reproductive strategy of field-collected hydra strains. We found no significant genetic change during the two years in the study population. Clone lines were detected between seasons and even years, suggesting that clonal lineages can persist for a long time in a natural population. We also found that distinct genotypes differ in sexual reproduction frequency, but these differences did not affect whether genotypes reappeared across samplings. Our study describes changes in population genetic structure across the seasons in a hydra population for the first time, providing key insights into the biology of the species, while also contributing to understanding the population biology of facultative sexual species inhabiting freshwater ecosystems.

## Introduction

Facultative sexual organisms, such as Cnidarians or Cladocerans are very important elements of marine and freshwater ecosystems, and their large numbers make them essential for the construction of aquatic food web system. Facultative sexual organisms often inhabit ephemeral or highly seasonal environments where favourable periods alternate with unfavourable ones in either predictable or unpredictable ways. In favourable periods, clonal reproduction often occurs, allowing the maximal utilization of available resources (Hadany & Otto, 2007; Stelzer, 2012; Stelzer & Lehtonen, 2016). On the other hand, the onset of adverse periods often trigger sexual reproduction, which ultimately results in the formation of resting eggs, such as in case of aphids (Simon et al., 2002) rotifers (Schröder, 2005; Stelzer & Lehtonen, 2016), water fleas (Tessier & Caceres, 2004) and hydras (Steele et al., 2019). The study of such reproductive systems has become more important nowadays, as the frequency of extreme conditions affecting habitats has significantly increased due to recent climate change. The importance of these studies is further reinforced by the large-scale disappearance or extinction of ecosystems (e.g. coral reefs, mangrove forests) engineered by facultative clonal species (Carpenter et al., 2008; Polidoro et al., 2010; Waycott et al., 2009). However, the population genetic structure of facultative sexual species can help them survive in highly variable environments, because different, sexually produced clonal lines respond differently to the environmental changes they face, while asexual reproduction allows quick growth of the populations. These enable such species to better adapt to the challenges posed by climate change and sustain high population sizes even under changing conditions (Pistevos et al., 2011).

The population genetic characteristics of facultative sexual organisms differ from those of obligate sexual organisms, partly because increase in their population size is not necessarily accompanied by the emergence of new genotypes, partly because selection affects them in different ways. For facultative sexual organisms, Nunney’s “lineage-selection” model suggests that solely asexual lines may enjoy short-term benefits (rapid exploitation of resources and high reproduction rates), but suffer disadvantages on the long term due to higher extinction rates compared to obligately sexual lines (Nunney, 1989). This lineage-selection model is probably even more important in a changing environment (e.g., seasonal habitats), as the creation of resistant formulas (e.g. resting eggs) is linked to sexual reproduction (e. g. in Daphnias, Decaestecker et al., 2009; in rotifers, Stelzer, 2012; or in hydras, Steele et al., 2019) and asexually reproducing lines can be easily removed by natural selection from the population when conditions deteriorate (Stelzer & Lehtonen, 2016). However, genotypes that reproduce only sexually can also be at a significant disadvantage in such an environment (Kokko, 2020), as they cannot reach a sufficiently large number of individuals during the optimal growth period. As a result, it is expected that in these populations genotypes that follow a strategy in which both modes of reproduction appear will prevail. In this case, the differences in the frequency and timing of their sexual reproduction (phenotypic plasticity of reproduction, Stelzer & Lehtonen, 2016), as has already been observed in rotifers (Tarazona et al., 2017), can greatly influence the survival of these genotypes. Thus, a special, seasonally changing genetic structure can emerge, in which the main driving force is the intermittent but large-scale genetic recombination due to clonal selection and sexual reproduction.

The genetic structure of populations of facultative sexual organisms, such as *Daphnia* species (cyclically parthenogenetic), is determined by the genetic consequences of combining sexual and asexual reproduction in the same life cycle (Carvalho, 1994; De Meester et al., 2006; Decaestecker et al., 2009), the pattern of which can be greatly influenced by the seasonal environment. At the start of the growing period (usually in spring), hatching of sexually produced dormant eggs increases the genetic variation in the population, between the offspring and hence the resultant unique clones (De Meester, 1996; De Meester et al., 2006). In contrast, asexual reproduction during the favourable season results in erosion of clone diversity through natural selection and extinction of clones, thus ultimately leading to lower genetic variation and deviations from Hardy-Weinberg equilibrium by the end of the favourable period (“clonal erosion”, De Meester, 1996; De Meester et al., 2006; Ortells et al., 2006; Tessier et al., 1992). Significant amounts of hidden genetic variation may be present in such populations that are not genetically expressed while the individuals are clonally reproducing but may be expressed after sexual recombination (Deng & Lynch, 1996; Pfrender & Lynch, 2000). Natural selection acts differently during the two reproductive phases (King & Schonfeld, 2001; Pfrender & Lynch, 2000), because during the asexual phase, all genes belong actually to one linked group, and so selection affects the whole genome. For this reason, clonal selection also shapes the interaction of genetic variation, in contrast to sexual reproduction, which interrupts the relationships of these linked alleles (Decaestecker et al., 2009). De Meester et al., (2006) identify three key factors (population size, length of the favourable season and strength of clonal selection) that influence the genetic structure of cyclical parthenogens. The extent to which “clonal erosion” affects the genetic structure of cyclic parthenogens is primarily determined by the three factors mentioned above (De Meester et al., 2006). The remarkable effect of clonal erosion on seasonal changes in the genetic composition of populations has already been shown by (Yin et al., 2010) in Daphnia, where the survival of clonal genotypes varied from population to population, significantly altering the genotype composition of the following year’s population. Furthermore, the picture is further complicated by the fact that life history strategies and physiological condition (e. g. age, body size, nutrition-state, stress-state) of individuals can also significantly influence genetic composition of populations (genotype diversity), as these are factors that clearly play a role in initiating sexual reproduction and thereby alter population dynamics (Hadany & Otto, 2007). However, the exact role of genotype-specific life history strategies have not been adequately explored in previous studies.

*Hydra oligactis* living in seasonal habitats of the temperate zone can serve as an excellent model animal for the ecological study of the reproductive system of facultative sexual organisms. *H. oligactis* polyps reproduce asexually (budding) throughout much of the year but switch to sexual reproduction in response to cooling (Reisa, 1973). In natural habitats, within the distribution range of *H. oligactis* sexual reproduction occurs from late summer to December (Ribi et al., 1985; Sebestyén et al., 2018; Welch & Loomis, 1924). During sexual reproduction, persistent/diapausing eggs are produced which tolerate desiccation and freezing (Steele et al., 2019). Based on our observations, however, in some of the natural populations, asexually and sexually reproducing individuals occur simultaneously before the unfavourable periods and adults can survive the unfavourable periods in large numbers (Miklós et al., 2021; MM & JT, personal observations). Therefore, the contribution of sexual individuals with diapausing eggs to the genetic structure and seasonal population dynamics of this species is unclear, as this has not been studied so far.

In our study we aimed at obtaining detailed genetic and phenotypic data of this model system thus to obtain more insight on the effect of genotype to reproductive strategies in this species. To this end, we genotyped hydra strains (using Restriction-site Associated DNA Sequencing, RAD-Seq) that were collected from a single population during spring and autumn in two years (four collections in total) and were kept under standard conditions in the laboratory to collect data on reproductive strategy. We used these data to ask firstly if clone lineages survive the unfavourable period and, if yes, how this affects the genetic composition of a population in such a seasonal environment in temperate climate. Secondly, we wanted to explore the role of different genotypes in the propensity for sexual reproduction and its timing, as well as in the associated population dynamics.

## Materials and methods

### Study design and field collection of Hydra polyps

Laboratory Hydra strains were established from a single oxbow lake in Eastern Hungary (Tiszadorogma, 47.6712 N, 20.8641 E) on four dates: 31st May 2018 (henceforth called Spring 2018), 1st October 2018 (Autumn 2018), 16th May 2019 (Spring 2019) and 24th September 2019 (Autumn 2019). A detailed description of sampling available in (Tökölyi et al., 2021).

Briefly, we collected polyps from multiple locations along the shoreline (distance between locations at least 2 meters). We recorded the GPS coordinates of each collection point (Garmin GPSMAP 60 CSx), (Fig. 1.; Supplementary Table 1). *Hydra* polyps were collected from free-floating and submerged macrophytes (most often *Ceratophyllum demersum, Ceratophyllum submersum, Myriophyllum spicatum, Stratiotes aloides*), then were put in a Falcon tube with lake water. On the day of collection, animals in Falcon tubes were transported to the laboratory in a cool box, where they were identified by stereomicroscopy (with Euromex StereoBlue stereo microscope) based on morphology — tentacle length / body length, the presence of stalk, tentacle formation in buds (Schuchert, 2010).

**Fig. 1.**
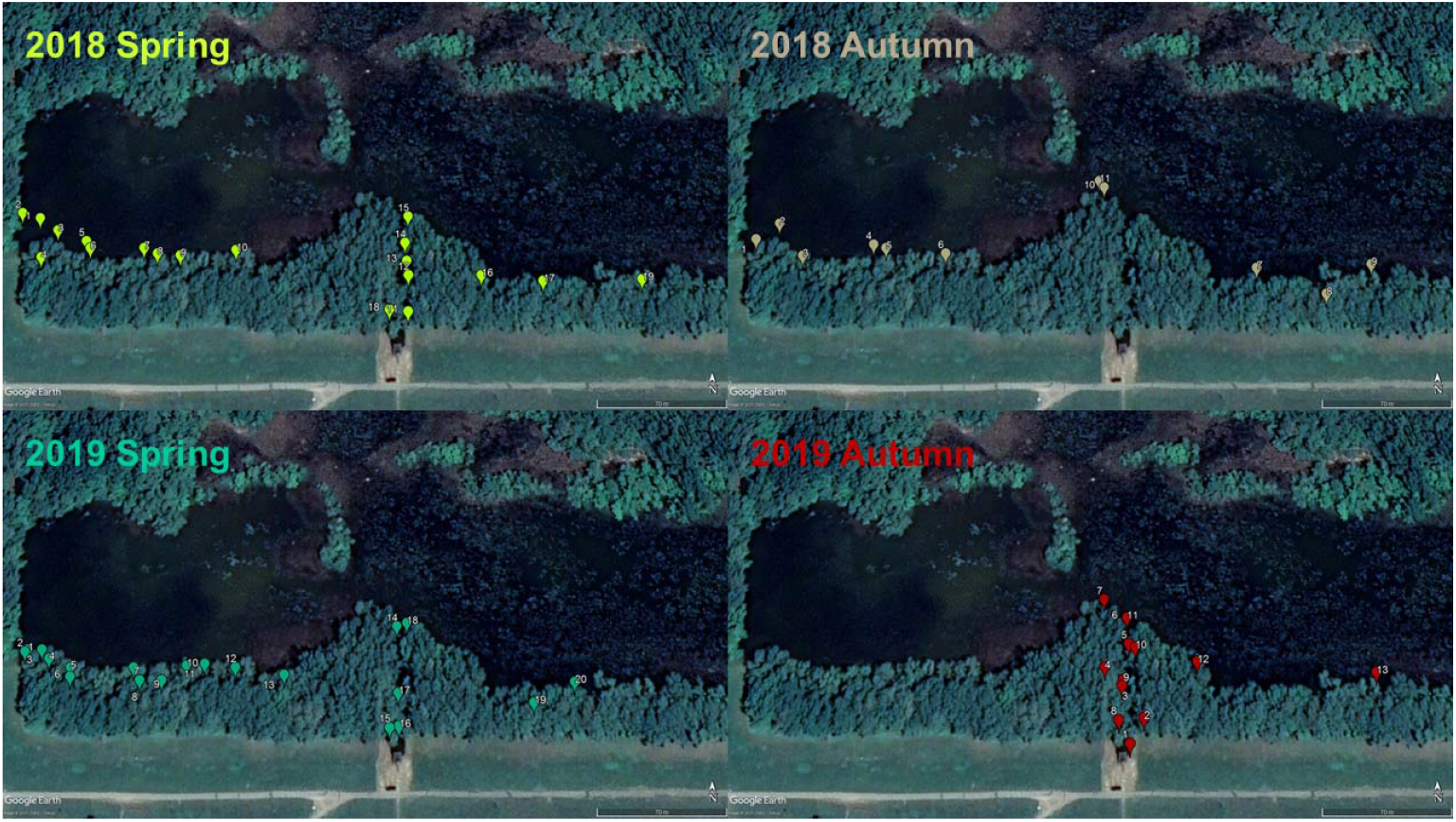
Map showing the collection point of H. oligactis polyps in two distinct seasons (spring vs. autumn) in two consecutive years (2018 and 2019, four samplings in total) from a single population in Central Hungary

### Lab maintenance of hydra strains and sex induction

Hydra polyps transported to the laboratory were immediately moved to hydra medium (M-solution: 1mM Tris, 1mM NaCl, 1mM CaCl2, 0.1mM KCl, 0.1mM MgSO4 at pH 7.6; Lenhoff, 1983). We selected up to five polyps from each location/collection to create strains from them through their budding (asexual reproduction). Both natural-collected polyps and their asexual offspring were kept individually in 6-well plastic plates, which contained 5 ml M-solution per well. Experimental animals were asexually propagated for 10 weeks in the first phase and then placed into cold circumstances to induce sexual reproduction in the second phase. Details of the standard living conditions of the hydra polyps, both for asexual reproduction phase and cooling phase, can be found in Tökölyi et al., (2021). Up to N=18 of polyps/strain were retained to collect data on reproductive mode. Experimental animals were kept for five months under second phase and were checked twice per week under a stereo microscope (with Euromex StereoBlue stereo microscope) to detect the start of gonadogenesis.

### Drying and DNA extraction

Asexual buds detached from experimental animals were used to genotype strains. They were dried using silica gel and stored at room temperature to preserve DNA quality (see Miklós et al., 2021). Genomic DNA from *H. oligactis* individuals was isolated using a standard mammalian nucleic acid extraction protocol (detailed description of used method, see in supplementary of Miklós et al., 2021; (Supplementary Methods 1)). The samples were stored in a freezer (−20 °C) until libraries were prepared.

### RAD-Seq library preparation

Details of the library preparation protocol can be found in the supplement of Miklós et al., 2021; (Supplementary Methods 1). Quality and quantity of the library were checked with Bioanalyzer (High-Sensitivity DNA Kit). Libraries were sequenced on an Illumina NovaSeq platform (paired-end, 150 nt) at NovoGene (Beijing, China). Our samples were sequenced in three separate RAD libraries (in the first 56 individuals, in the second 53 individuals and in the third 30 individuals). In all library preparations, we used the same methodologies, but in the third library, only 15 cycles were used during PCR amplification, to reduce the presence of PCR duplicates. Finally, GenBank association numbers associated with identified specific genotypes were recorded (Supplementary Table 1).

### Sequence processing and decontamination

Firstly, raw Illumina reads were processed using Stacks *process_radtags* pipeline (Catchen et al., 2013; Miklós et al., 2021). We first ran the pipeline with default parameters and calculated the GC content of the resulting RAD loci with the BBmap suite of tools (https://sourceforge.net/projects/bbmap/). This showed a secondary GC peak suggesting sequence contamination from bacterial DNA. Therefore, we performed *in silico* decontamination using the NCBI Basic Alignment Search Tool (BLAST, v. 2.7.1; Altschul et al., 1990) to map sequences to the NCBI nucleotide collection database (*nt*, downloaded 31^st^ March 2021), with *blastn* task and E-value cutoff set to 1e-05. RAD loci whose best match was a cnidarian sequence in the *nt* database or showed no hit were retained, while loci that mapped to any other taxonomic group were separated to form a contaminants database. We then mapped our paired-end demultiplexed reads to this contaminants database with Bowtie 2 *--very sensitive* and retained only unmapped reads. Sequence handling was done with the BBmap suite of tools (https://sourceforge.net/projects/bbmap/), while taxonomic annotation was done with the *taxonomizr* R package (v. 0.5.3; Sherrill-Mix, 2019); R Core Team, 2020).

Next, we ran the *de novo* pipeline on the unmapped reads, with the following settings: minimum depth of coverage required to create a stack (-m) of 3, the number of mismatches allowed between stacks within individuals (-M) of 3 and the number of mismatches allowed between stacks between individuals (-n) of 3. These parameters were selected based on our previous study (Miklós et al., 2021) that included some individuals from the population now being studied.

From the resulting set of loci, we retained those that were shared by at least 80% of the sample. Additional filtering was performed on the resulting locus catalog using VCFtools (Danecek et al., 2011), with the following parameters: we required a minor allele count (--mac) of 3, minor allele frequency (--maf) of 0.05, minimum genotype quality (--minGQ) of 30, minimum read depth (--minDP) of 10 and mean depth values across individuals (--max-meanDP) less than 97.5% of the mean depth values for the whole dataset. Then, we checked individual missingness, removed samples with >50% missingness and repeated all filtering steps without these samples.

### Clone detection and sibship reconstruction

To identify clones, we first inspected the spectrum of genetic diversity, i.e. the distribution of pairwise genetic distances of the samples (Rozenfeld et al., 2007). Clonally derived individuals in theory should be genetically identical to each other. However, due to sequencing errors and somatic mutations they frequently show a distribution of genetic distances >0, but less then the genetic distance of distinct genotypes, which often results in a distinct peak of low genetic distances on the histogram. We also used the software COLONY (v. 2.0.6.6; Jones & Wang, 2010) to infer clones more formally using an optimized threshold that takes into account mistyping rates, missing data and the number and allele frequencies of markers (Wang, 2016). The method implemented in COLONY uses a likelihood framework to assign individuals to candidate relationships of clone mates and close competitive relationships (e.g. full sibships) and has been shown to accurately identify individuals belonging to the same multilocus genotypes (MLGs) through simulations (Wang, 2016).

All individuals were included as potential offspring in the analysis, since the presence of clonality in *Hydra* implies that generations can be overlapping and there is no unequivocal way to assign candidate parents. These potential offspring were then assigned into clonal lineages. For the COLONY analysis, we used a full-likelihood-pair-likelihood score combined (FPLS) method, assumed a polygamous mating system for both parents and kept all other parameters at their default values. Initial error rates were set to 0.01 for both allelic dropout rate and other error rate of each locus in COLONY.

### Seasonal genetic structure

To visualize the seasonal distribution of sample genotypes, minimum spanning networks (MSN) were constructed using the function *poppr*.*msn* (Kamvar et al., 2014) in R. The network was constructed on the basis of genetic distance matrix calculated in *ape*’s R package *dist*.*gene* function, with pairwise deletion of missing loci. These relationships were visualized with MSN (because for clonal organizations it can be a better visualization tool than tree drawing methods) generated using the R packages *igraph* and *poppr* (Csardi & Nepusz, 2005; Kamvar et al., 2014).

Basic population genetic statistics (expected heterozygosity, observed heterozygosity, fixation index, allelic richness, the number of private alleles and clonal richness, evenness and diversity) were calculated for two datasets. In the first analysis, all samples were included. However, since *H. oligactis* is a clonal species, the presence of clones can bias the results of population genetics statistics. Therefore, we also prepared a reduced dataset that included one strain from each MLG per sampling (based on results obtained from COLONY) and repeated the calculations on this reduced dataset.

Expected heterozygosity, observed heterozygosity and fixation index were calculated in the *hierfstat* R package (v. 0.5-7; Goudet et al., 2020), allelic richness was calculated with the *PopGenReport* R package (v. 3.0.4; Adamack & Gruber, 2014), while the number of private alleles were obtained with the *poppr* R package (v. 2.9.0; Kamvar et al., 2014). Clonal richness was calculated as R = (G-1) / (N-1), where G is the number of MLGs and N is the number of strains (Dorken & Eckert, 2001). Evenness and Shannon-Wiener diversity were calculated in *poppr*.

To detect genetic structure with respect to sampling dates, we performed discriminant analysis of principal components (DAPC; Jombart et al., 2010) on the reduced dataset. DAPC analysis was performed using the *adegenet* (v. 2.1.1) package in R (Jombart, 2008; R Core Team, 2020). The number of principal components used in the DAPC analysis was set to 13 following alpha-score optimization and we generated inertia ellipses encompassing ∼67% of the cloud of points for each population. For the DAPC analysis we only included 1 individual from each MLG.

Finally, we performed Analysis of Molecular Variance (AMOVA) between seasons. AMOVA was implemented in *R’s pegas* (v. 0.10) package (Paradis, 2010) using 1000 permutations.

## Results

### Establishment of field-collected strains in laboratory

We established strains from N = 211 polyps (N = 54, 40, 59, 58, respectively for the four collection dates). However, some of these strains were lost due to mortality before yielding usable data and DNA samples. A total of 138 strains were genotyped (N = 40, 30, 38, 30 for the four seasons).

### Read statistics and decontamination

There were altogether 699.8 million raw paired-end reads. From these raw reads, 93.6% were retained after filtering for low-quality reads, adapter contamination, ambiguous barcodes and ambiguous RAD-tags. There were an average of 4.7 million reads per sample (range 2.3-20.9 million).

Running the Stacks *de novo* pipeline with default settings identified 1 602 803 million RAD loci. 46.9% percent of these loci showed no hit in the *nt* database, a further 13.6% percent mapped to cnidarian sequences. The remainder mapped to other taxonomic groups and were filtered out to form a contaminants database. The top contaminants were Pseudomonadales and Burkholderiales (Supplementary Fig 1.), two bacterial orders that are commonly found within the *Hydra* microbiome (Fraune et al., 2015). After removing presumed contaminant loci, the secondary GC peak diminished substantially (Supplementary Fig. 1).

The proportion of read pairs identified to be PCR duplicates after removal of contaminants was 27.2%. Mean coverage after removing contaminants and PCR duplicates was 13.7x per sample (range: 4.1x - 25.7x).

Running Stacks *de novo* with these final settings, we obtained 1417 RAD loci after filtering. Six samples (all of them from the 2019 spring collection) had individual missingness >50% and were removed. Repeating the filtering of loci without this sample resulted in a final set of 2257 loci. Individual missingness was 15.2% (range: 2.3-54.1%).

### Clone detection

Inspection of the spectrum of genetic diversity showed no clear threshold to delineate multilocus genotypes (Supplementary Figure 2). While there was a clear peak of low genetic distance (less than ∼0.06), we also detected a smaller, secondary peak at genetic distance ∼0.11. This secondary peak could stem from the fact that genotyping error rates are higher in some of the samples (as seen in our previous study for samples with low coverage; Miklós et al., 2021), or because somatic mutations are more prevalent in some of the samples. Given that we used very stringent filtering of SNPs (minimum genotype quality of 30 and minimum read depth of 10), we think that genotyping error rates in general should be low. Nonetheless, the presence of this secondary peak in the spectrum of genetic diversity makes the identification of clones more difficult.

The COLONY analysis also revealed these difficulties. We identified N=53 multilocus genotypes in the set of N=132 strains included in the analysis (Fig. 2). However, 11 of these 53 MLGs (each of them consisting of a single strain) were inferred with a probability <0.9 (while all other MLGs were inferred with probability 1.0). Therefore, we decided to remove these individuals from subsequent analyses since we cannot unequivocally assign them to MLGs. As a consequence, subsequent analyses are presented for N=121 strains belonging to 42 MLGs. Most of these MLGs (N=35, 83.3%) were detected in only one season. Five MLGs (11.9%) were detected in two seasons, one (2.4%) in three seasons and one (2.4%) in all four seasons.

**Fig 2.**
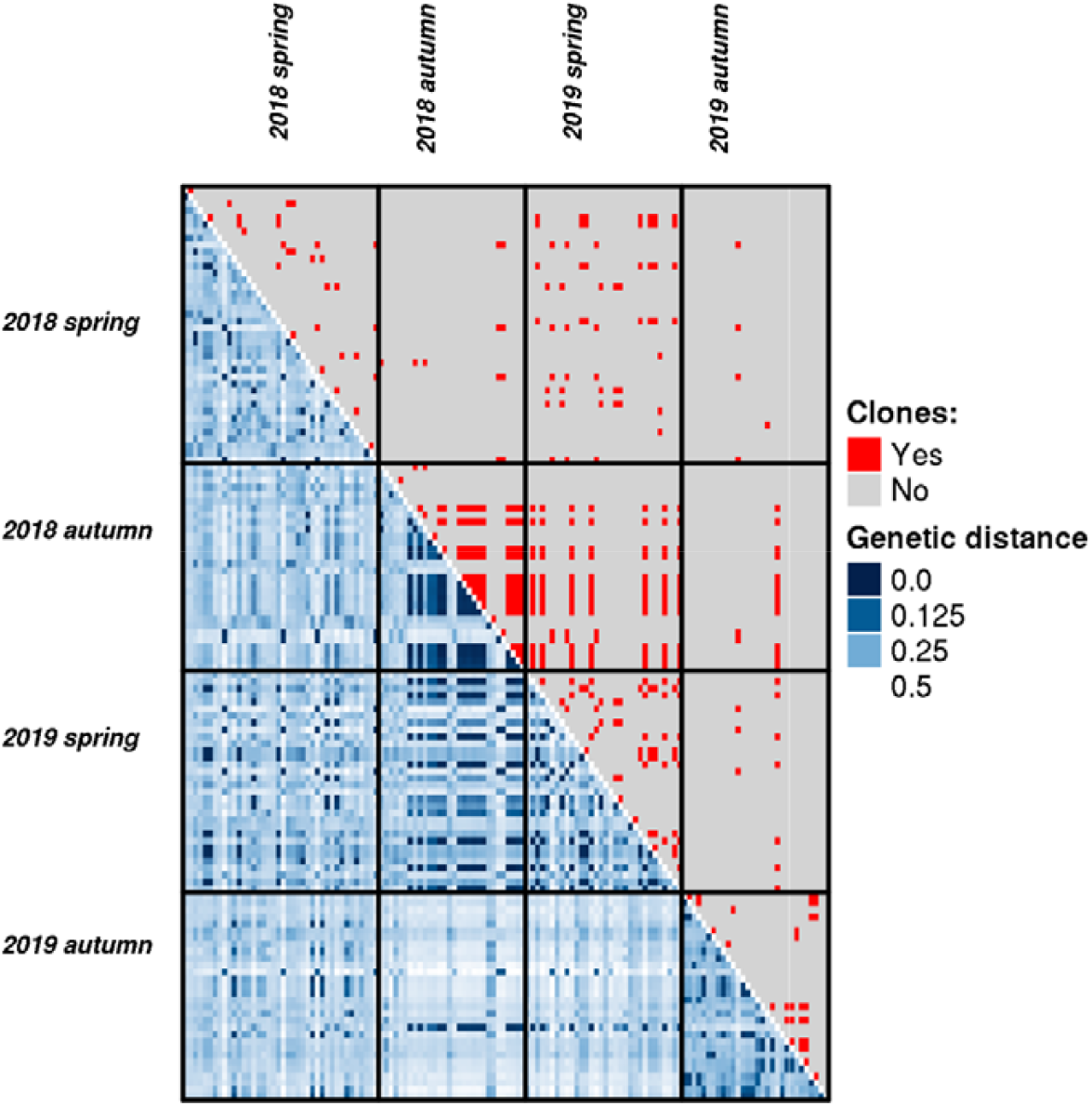
Heatmap showing genetic distance between N=121 genotyped *Hydra oligactis* strains collected at four distinct time points (lower diagonal). Pairs of strains that were inferred to be clones in the COLONY analysis are shown in red in the upper diagonal.

### Seasonal genetic structure

Clonal richness and evenness did not show clear seasonal trends, although Shannon-Wiener diversity was somewhat higher in the spring samples. Observed heterozygosity, expected heterozygosity, F_is_, allelic richness and the number of private alleles did also not show marked seasonal trends (Table 1.).

**Table 1.** Basic population genetic statistics from 2557 SNPs obtained with RAD-Seq for N=121 *H. oligactis* strains established from polyps collected in two distinct seasons (spring vs. autumn) in two consecutive years (2018 and 2019, four samplings in total) from a single population in Central Hungary.

| Full dataset (N=121) |  |  |  |  |  |  |  |  |  |  |  | Clone-corrected dataset (N=54) |  |  |  |  |
| --- | --- | --- | --- | --- | --- | --- | --- | --- | --- | --- | --- | --- | --- | --- | --- | --- |
| Year | Season | N | MLGs | Clonal<br>richness | Clonal<br>Evenness | Shannon-Weiner<br>diversity | H<sub>obs</sub> | H<sub>exp</sub> | F<sub>is</sub> | Allelic<br>richness | Private<br>alleles | H<sub>obs</sub> | H<sub>exp</sub> | F<sub>is</sub> | Allelic<br>richness | Private<br>alleles |
| 2018 | Spring | 39 | 21 | 0.53 | 0.85 | 2.88 | 0.31 | 0.24 | -0.26 | 1.78 | 1 | 0.30 | 0.23 | -0.24 | 1.44 | 5 |
| 2018 | Autumn | 30 | 11 | 0.34 | 0.54 | 1.87 | 0.34 | 0.24 | -0.41 | 1.71 | 0 | 0.35 | 0.26 | -0.31 | 1.48 | 1 |
| 2019 | Spring | 31 | 12 | 0.37 | 0.76 | 2.14 | 0.33 | 0.24 | -0.36 | 1.74 | 0 | 0.34 | 0.25 | -0.29 | 1.47 | 0 |
| 2019 | Autumn | 21 | 8 | 0.35 | 0.82 | 1.86 | 0.38 | 0.26 | -0.41 | 1.75 | 8 | 0.37 | 0.26 | -0.34 | 1.50 | 9 |

The Minimum Spanning Network showed that individuals collected from different seasons did not show significant separation from each other, but clustered into 3 major branches regardless of the time of collection (Fig. 3.).

**Fig. 3.**
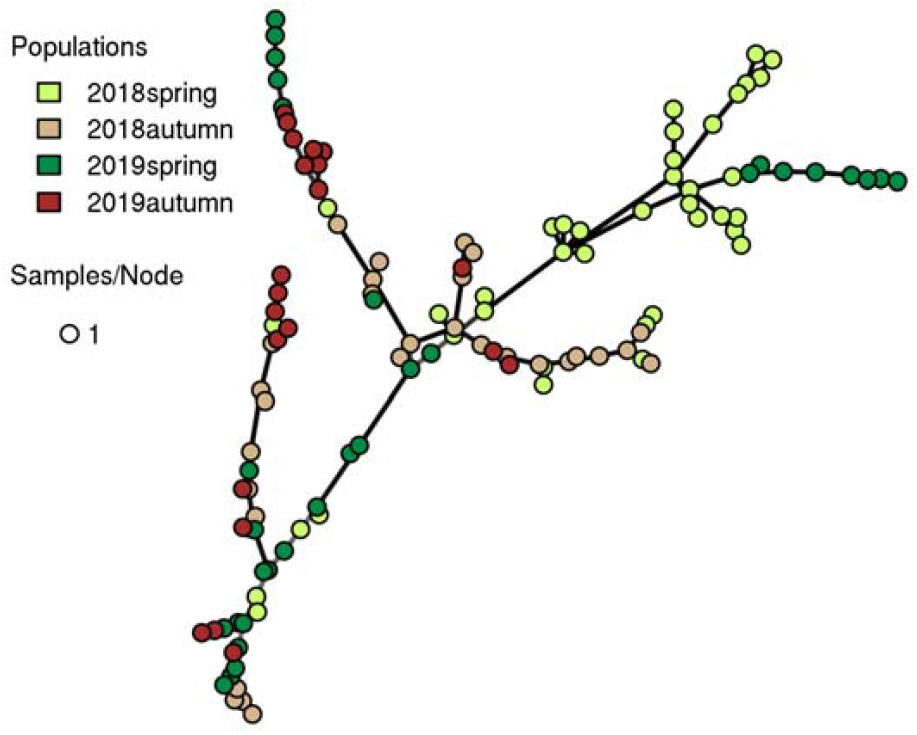
Minimum spanning network of N=121 *H. oligactis* strains established from polyps collected in two distinct seasons (spring vs. autumn) in two consecutive years (2018 and 2019, four samplings in total) from a single population in Eastern Hungary.

The DAPC analysis likewise indicated substantial overlap between seasons, with the final sample being more distinct from the rest (Fig. 4). Reflecting this fact, we found significantly different genetic structure between seasons based on AMOVA (σ^2^ = 0.003, p=0.01).

**Fig. 4.**
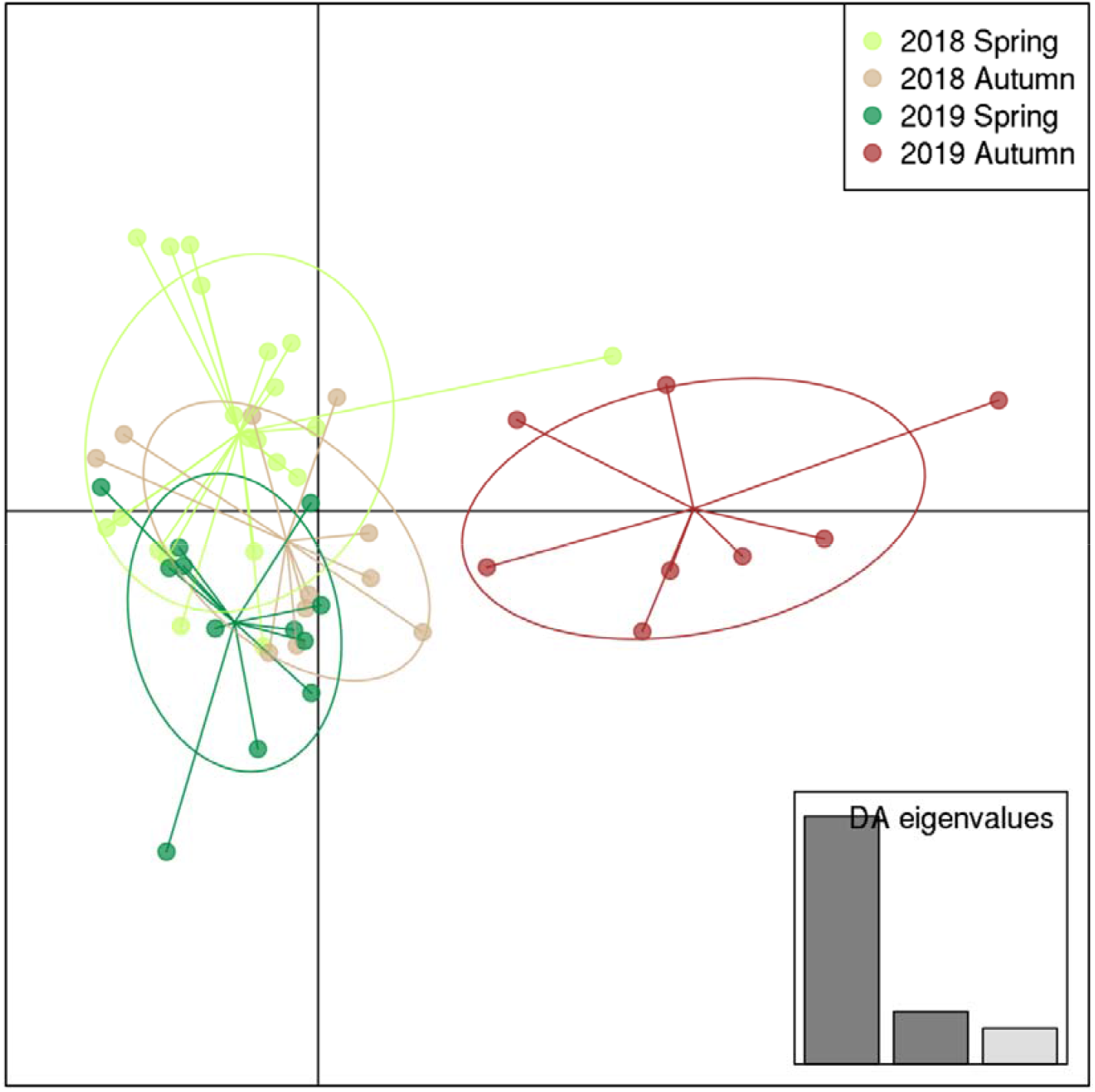
Differentiation of strains derived from four distinct sampling types based on Discriminant Analysis of Principal Components (DAPC) performed on a reduced dataset containing one individual from each MLG per sampling.

### Genetic structure and reproductive mode

Reproductive mode was inferred based on data from N = 921 polyps (this is a subset of individuals analysed in Tökölyi et al., (2021) that belong to the 121 genotyped strains). There were on average 7.74 polyps per strain to estimate reproductive mode (range: 1-18). We could not estimate reproductive mode for two strains where all polyps were lost before data collection. After genotyping, there were 21.9 polyps per MLG to estimate reproductive mode (range: 2-136).

With the exception of one fully asexual MLG, all MLGs contained a mixture of sexual and asexual individuals. The average proportion of sexual individuals per MLG was 74.0% and 11 out of 42 MLGs (26.2%) contained only sexual individuals. The proportion of variance in reproductive mode explained by MLG ID based on the binomial GLMM was 46.7%, while season and polyp age explained 10.1% of variance in total.

There was no difference in the proportion of sexual individuals in MLGs that were observed in multiple seasons compared to those that were observed only in a single season (Kruskal test, χ^2^ = 0.014, p = 0.905).

## Discussion

The primary objective of our study was to investigate how the genetic composition of a population of a common freshwater facultative sexual organism, *Hydra oligactis*, changes in a temperate seasonal environment. We detected (1) limited changes in the seasonal population genetic composition and found that: 2) some hydra clone lines can survive between years and seasons. Furthermore, we also found that 3) distinct genotypes differ in sexual reproduction frequency. Finally (4) the above differences did not affect whether these genotypes reappear between samplings. We discuss the consequences and circumstances of these findings below.

We found no clear evidence for an abundance of spring genotypes in the Hydra population, despite the fact that it would be a logical consequence of their high rate of sexual reproduction in winter (generation of new genotypes) and has been described previously for other facultative sexual organisms (Daphnia; De Meester, 1996; De Meester et al., 2006). Based on previous studies, we would have expected that asexual reproduction during the favourable period would lead to a reduction in clonal diversity (through natural selection and random extinction of clones), resulting in lower genetic variation at the end of the favourable period and greater deviations from Hardy-Weinberg equilibrium (clonal erosion; De Meester, 1996; De Meester et al., 2006; Tessier et al., 1992), but in our study hydra population, we were not able to detect this clonal erosion clearly. One simple reason for this could be that we did not include enough individuals in the study and thus stochastic effects due to random sampling could affect our results. Another reason could be that we could not actually identify all clones when analysing the data. Identifying clones in this study proved difficult because of the relatively high frequency of pairs of individuals that had a genetic distance intermediate between clones and distinct genotypes. We used very stringent filtering of SNPs to increase genotyping accuracy, which further reduced sample size. However, all retained strains were assigned to MLGs with a high certainty in the COLONY analysis, therefore we think that the obtained results reflect true biological patterns. In general, we can conclude that there is definitely a set of genotypes in the population that persists for a relatively long time regardless of seasonality. In addition, there was noticeable a major change in genotype composition by the second autumn, but the exact reason for this is not known (it is possible that this was a consequence of an unusually warm summer before, which may have enhanced and modified the usual selection effects).

This facultative sexual reproduction system has several implications for the genetic structure and evolution of such populations. One is that through selection, genotypes with weak competitive ability disappear from the population, so only surviving genotypes can trigger sexual reproduction. Another effect is that different hidden genetic variations are not expressed during asexual reproduction, but which may be expressed after sexual recombination (Deng & Lynch, 1996; Pfrender & Lynch, 2000), thus allowing populations to develop new traits increasing their adaptability. In addition, it should be mentioned that spring genotype can also contribute to abundance through gene flow from individuals from other populations and persistent eggs, as evidenced by our previous study (Miklós et al., 2021). However, we do not yet have definitive information on the mechanism, frequency, and time distribution of migration between these populations. Probable propagation vectors may be waterfowl, whose larger-scale migration occurs primarily in spring and autumn in this climate, so it is likely that gene flow may also be significant during these periods. Conversely, whether individuals migrate in either autumn or spring, they may contribute to genotype abundance primarily in spring, as individuals entering spring are equally affected by selection during the growth phase and individuals entering in autumn are much less likely to survive and reproduce in the new population. Overall, we were able to show that the effect of natural selection and the survival of certain genotypes without sexual reproduction are important factors in forming the genetic composition of facultative sexual hydra populations.

We reliably demonstrated that some clonal lines can survive seasons and even years, contrary to the literature that says that hydra polyps die in winter due to freezing waters, and only survive in sexual enduring forms (Brien, 1953). A survival strategy based on persistent eggs of sexual reproduction is common in aquatic invertebrate species that reproduce facultatively in temperate or cold climates, such as Rotifers (Walsh et al., 2014) or Cladocerans (Decaestecker et al., 2009). In climates where mild winters often occur, the survival of clones can also have a significant adaptive advantage in terms of a given clone genotype, resulting in a better return on investment in winter asexual reproduction. It has already been observed in Cladocerans that greater investment in asexual reproduction may be adaptive in a relatively mild winter climate where the risk of freezing is small in winter and the adaptive value of dormancy may be low (Tessier & Caceres, 2004) The situation may be similar for other temperate species, such as hydras. From a population dynamics point of view, such surviving individuals can significantly influence the composition of the early spring population, as they may even begin their clonal reproduction while the other genetic lines are still waiting in resting form. Thus, such hydra lines may start with a larger number of individuals over the newly emerging genotypes, thus gaining a significant advantage in resource exploitation. Nonetheless, our data set collected over two years suggests that this survival may not be of such an advantage to such lines that most new genotypes can be displaced but may significantly contribute to the persistent presence of these genotypes in the population.

We also observed that different genotypes differ in sexual reproduction frequency. Based on few previous studies (Tökölyi et al., 2017; Tomczyk et al., 2015), genotypic differences may contribute to variation in the propensity of sexual reproduction in hydras because, under normal laboratory conditions, *H. oligactis* strains express differences in the probability of initiating sexual reproduction and in post-sexual survival rates (Ngo et al., 2021). Conversely, in another previous study analysing reproductive mode under field condition, we detected a high rate of phenotypic plasticity in reproduction strategy in this species (Miklós et al., 2021). Furthermore, this also raises the possibility of a genotype responding differently to the same environmental stimulus (even depending on its internal condition; Hadany & Otto, 2007) thus creating a diverse reproductive state in the populations depending on the season (Tökölyi et al., 2021). Our previous statement was confirmed in this study however, we also showed that different genotypes show a different willingness to initiate sexual reproduction under the same conditions. This may play a significant role in the long-term change in the genetic composition of such populations, as the genotypes best adapted to a given environmental condition can be the most widespread in a given habitat. Moreover, such a facultative sexual reproductive system is likely to allow rapid adaptation to local conditions through selection of genotypes with an adaptive combination of phenotypic plasticity responses (De Meester et al., 2004). Some studies have already provided evidence for effectual tracking of environmental changes over time in natural Daphnia populations (Cousyn et al., 2001; Hairston et al., 2001). That is, in mild climates where the winter is less harsh, genotypes that show a lower propensity to initiate sexual reproduction are preferred. In addition, in previous cases, relevant differences between clone lines (genetic individuals) in some marine species in their response to environmental stimuli, including the choice of mode of reproduction, have been described (Langer et al., 2009; Pistevos et al., 2011). This trait is of great importance for these species in adapting to the environment without further mutational novelties, driven only by selection. This is also important because such metazoan will be more easily able to adapt to rapid global change through natural selection affecting existing genotypic variations (Balanyá et al., 2006; Bradshaw & Holzapfel, 2001), despite the slow onset of their mutational changes (Hoffmann et al., 2003).

Interestingly, differences in genotype traits did not affect the reappearance of specific genotypes in different samples. The simplest explanation for this may be that such a sampling time interval is not sufficient to accurately describe such consequences of population dynamic effects, because random effects may still obscure them. An alternative explanation could be that each genotype is so plastic that even if a significant proportion of individuals in genetic lineages are likely to reproduce sexually, they may also be survived by asexually reproducing polyps, which thus maintain the genetic lineage (bet-hedging; Simons, 2009; Steele et al., 2019).

## Conclusion

Overall, we have been able to show limited variation in the genetic composition of the seasonal population, and that some hydra clonal lines can survive between years and seasons. We also observed that distinct genotypes differ in the frequency of sexual reproduction. Finally, we found that these observed differences did not affect whether these genotypes would reappear among samples. In conclusion, these findings suggest that facultative sexual organisms are genetically plastic enough to maintain different reproductive strategies (asexual and asexual reproduction) in parallel, which may give them a significant advantage in predictably changing environments, thus increasing their adaptive capacity even in the face of unpredictable changes. This makes the study of this ability even more relevant today, as ecosystems with populations with these traits could be the key to mitigating ecological damage caused by climate change.

## Acknowledgements

We are grateful to Jinliang Wang for help with the COLONY analysis. This study was supported by NKFIH grant FK 124164. ZB was supported by the Thematic Excellence Programme (TKP2020-IKA-04) of the Ministry for Innovation and Technology in Hungary. Finally, we are thankful to the following people for lab assistance: Réka Gergely, Berta R-Almási, Beatrix Kozma, Dávid Tenkei, Erzsébet Ágnes Nehéz, and Flóra Sebestyén.

## Data Availability Statement

Raw DNA sequence reads have been deposited in the U.S. National Center for Biotechnology Information (NCBI) Sequence Read Archive (Accession number SUB10367137). The R-scripts of our analysis was uploaded to Github (https://github.com/jtokolyi/Hydra_oligactis_PopDynamics).

## Author Contributions

M.M., L. L., and G. S. performed molecular lab work. M.M., and J.T. collected samples. M.M., J.T. and Z.B. performed analyses. All authors contributed to writing the manuscript.

