## Supplementary Table 1, Supplementary Fig 1.,Supplementary Fig 2. for "Seasonal variation of genotypes and reproductive plasticity in a facultative clonal freshwater invertebrate animal (Hydra oligactis) living in a temperate lake"

Supplementary Tables:

Supplementary Table 1. Location IDs, Collection dates, GPS coordinates, Sampling occasions, GenBank association numbers of the collection sites of sequenced *H. oligactis* polyps.

| **Individual ID** | **Location ID** | **Collection Date** | **Sampling occasion** | **GPS cordinates** | **GenBank association number** |
| --- | --- | --- | --- | --- | --- |
| M28_2018osz_1_3 | M28/1 | 2018.10.01 | 2018 Autumn | N47.67121 E20.86296 | SAMN21437818 |
| M28_2018osz_2_1 | M28/2 | 2018.10.01 | 2018 Autumn | N47.67130 E20.86312 | SAMN21437819 |
| M28_2018osz_2_2 | M28/2 | 2018.10.01 | 2018 Autumn | N47.67130 E20.86312 | SAMN21437820 |
| M28_2018osz_2_3 | M28/2 | 2018.10.01 | 2018 Autumn | N47.67130 E20.86312 | SAMN21437821 |
| M28_2018osz_3_3 | M28/3 | 2018.10.01 | 2018 Autumn | N47.67112 E20.86335 | SAMN21437822 |
| M28_2018osz_3_4 | M28/3 | 2018.10.01 | 2018 Autumn | N47.67112 E20.86335 | SAMN21437823 |
| M28_2018osz_4_1 | M28/4 | 2018.10.01 | 2018 Autumn | N47.67118 E20.86389 | SAMN21437824 |
| M28_2018osz_4_4 | M28/4 | 2018.10.01 | 2018 Autumn | N47.67118 E20.86389 | SAMN21437825 |
| M28_2018osz_5_1 | M28/5 | 2018.10.01 | 2018 Autumn | N47.67116 E20.86399 | SAMN21437826 |
| M28_2018osz_5_2 | M28/5 | 2018.10.01 | 2018 Autumn | N47.67116 E20.86399 | SAMN21437827 |
| M28_2018osz_5_3 | M28/5 | 2018.10.01 | 2018 Autumn | N47.67116 E20.86399 | SAMN21437828 |
| M28_2018osz_5_4 | M28/5 | 2018.10.01 | 2018 Autumn | N47.67116 E20.86399 | SAMN21437829 |
| M28_2018osz_6_1 | M28/6 | 2018.10.01 | 2018 Autumn | N47.67113 E20.86446 | SAMN21437830 |
| M28_2018osz_6_2 | M28/6 | 2018.10.01 | 2018 Autumn | N47.67113 E20.86446 | SAMN21437831 |
| M28_2018osz_6_3 | M28/6 | 2018.10.01 | 2018 Autumn | N47.67113 E20.86446 | SAMN21437832 |
| M28_2018osz_6_4 | M28/6 | 2018.10.01 | 2018 Autumn | N47.67113 E20.86446 | SAMN21437833 |
| M28_2018osz_7_1 | M28/7 | 2018.10.01 | 2018 Autumn | N47.67105 E20.86687 | SAMN21437834 |
| M28_2018osz_7_2 | M28/7 | 2018.10.01 | 2018 Autumn | N47.67105 E20.86687 | SAMN21437835 |
| M28_2018osz_7_3 | M28/7 | 2018.10.01 | 2018 Autumn | N47.67105 E20.86687 | SAMN21437836 |
| M28_2018osz_7_4 | M28/7 | 2018.10.01 | 2018 Autumn | N47.67105 E20.86687 | SAMN21437837 |
| M28_2018osz_8_2 | M28/8 | 2018.10.01 | 2018 Autumn | N47.67091 E20.86738 | SAMN21437838 |
| M28_2018osz_8_3 | M28/8 | 2018.10.01 | 2018 Autumn | N47.67091 E20.86738 | SAMN21437839 |
| M28_2018osz_9_1 | M28/9 | 2018.10.01 | 2018 Autumn | N47.67107 E20.86776 | SAMN21437840 |
| M28_2018osz_9_3 | M28/9 | 2018.10.01 | 2018 Autumn | N47.67107 E20.86776 | SAMN21437841 |
| M28_2018osz_10_1 | M28/10 | 2018.10.01 | 2018 Autumn | N47.67151 E20.86570 | SAMN21437842 |
| M28_2018osz_10_3 | M28/10 | 2018.10.01 | 2018 Autumn | N47.67151 E20.86570 | SAMN21437843 |
| M28_2018osz_10_4 | M28/10 | 2018.10.01 | 2018 Autumn | N47.67151 E20.86570 | SAMN21437844 |
| M28_2018osz11_1 | M28/11 | 2018.10.01 | 2018 Autumn | N47.67155 E20.86566 | SAMN21437845 |
| M28_2018osz_11_2 | M28/11 | 2018.10.01 | 2018 Autumn | N47.67155 E20.86566 | SAMN21437846 |
| M28_2018osz_11_3 | M28/11 | 2018.10.01 | 2018 Autumn | N47.67155 E20.86566 | SAMN21437847 |
| M28_2018osz_11_4 | M28/11 | 2018.10.01 | 2018 Autumn | N47.67155 E20.86566 | SAMN21437848 |
| M28_2018tav_1_1 | M28/1 | 2018.05.31 | 2018 Spring | N47.67133 E20.86300 | SAMN21437849 |
| M28_2018tav_1_2 | M28/1 | 2018.05.31 | 2018 Spring | N47.67133 E20.86300 | SAMN21437850 |
| M28_2018tav_1_3 | M28/1 | 2018.05.31 | 2018 Spring | N47.67133 E20.86300 | SAMN21437851 |
| M28_2018tav_2_1 | M28/2 | 2018.05.31 | 2018 Spring | N47.67136 E20.86285 | SAMN21437852 |
| M28_2018tav_2_2 | M28/2 | 2018.05.31 | 2018 Spring | N47.67136 E20.86285 | SAMN21437853 |
| M28_2018tav_3_1 | M28/3 | 2018.05.31 | 2018 Spring | N47.67126 E20.86316 | SAMN21437854 |
| M28_2018tav_3_4 | M28/3 | 2018.05.31 | 2018 Spring | N47.67126 E20.86316 | SAMN21437855 |
| M28_2018tav_4_2 | M28/4 | 2018.05.31 | 2018 Spring | N47.67111 E20.86307 | SAMN21437856 |
| M28_2018tav_4_3 | M28/4 | 2018.05.31 | 2018 Spring | N47.67111 E20.86307 | SAMN21437857 |
| M28_2018tav_5_2 | M28/5 | 2018.05.31 | 2018 Spring | N47.67120 E20.86340 | SAMN21437858 |
| M28_2018tav_5_3 | M28/5 | 2018.05.31 | 2018 Spring | N47.67120 E20.86340 | SAMN21437859 |
| M28_2018tav_6_1 | M28/6 | 2018.05.31 | 2018 Spring | N47.67116 E20.86344 | SAMN21437860 |
| M28_2018tav_6_2 | M28/6 | 2018.05.31 | 2018 Spring | N47.67116 E20.86344 | SAMN21437861 |
| M28_2018tav_6_3 | M28/6 | 2018.05.31 | 2018 Spring | N47.67116 E20.86344 | SAMN21437862 |
| M28_2018tav_7_1 | M28/7 | 2018.05.31 | 2018 Spring | N47.67116 E20.86386 | SAMN21437863 |
| M28_2018tav_7_2 | M28/7 | 2018.05.31 | 2018 Spring | N47.67116 E20.86386 | SAMN21437864 |
| M28_2018tav_7_3 | M28/7 | 2018.05.31 | 2018 Spring | N47.67116 E20.86386 | SAMN21437865 |
| M28_2018tav_8_2 | M28/8 | 2018.05.31 | 2018 Spring | N47.67113 E20.86397 | SAMN21437866 |
| M28_2018tav_8_3 | M28/8 | 2018.05.31 | 2018 Spring | N47.67113 E20.86397 | SAMN21437867 |
| M28_2018tav_9_1 | M28/9 | 2018.05.31 | 2018 Spring | N47.67112 E20.86415 | SAMN21437868 |
| M28_2018tav_9_2 | M28/9 | 2018.05.31 | 2018 Spring | N47.67112 E20.86415 | SAMN21437869 |
| M28_2018tav_9_3 | M28/9 | 2018.05.31 | 2018 Spring | N47.67112 E20.86415 | SAMN21437870 |
| M28_2018tav_10_2 | M28/10 | 2018.05.31 | 2018 Spring | N47.67115 E20.86458 | SAMN21437871 |
| M28_2018tav_10_3 | M28/10 | 2018.05.31 | 2018 Spring | N47.67115 E20.86458 | SAMN21437872 |
| M28_2018tav_11_1 | M28/11 | 2018.05.31 | 2018 Spring | N47.67082 E20.86591 | SAMN21437873 |
| M28_2018tav_12_1 | M28/12 | 2018.05.31 | 2018 Spring | N47.67101 E20.86592 | SAMN21437874 |
| M28_2018tav_13_1 | M28/13 | 2018.05.31 | 2018 Spring | N47.67109 E20.86591 | SAMN21437875 |
| M28_2018tav_13_2 | M28/13 | 2018.05.31 | 2018 Spring | N47.67109 E20.86591 | SAMN21437876 |
| M28_2018tav_14_1 | M28/14 | 2018.05.31 | 2018 Spring | N47.67119 E20.86590 | SAMN21437877 |
| M28_2018tav_14_2 | M28/14 | 2018.05.31 | 2018 Spring | N47.67119 E20.86590 | SAMN21437878 |
| M28_2018tav_14_3 | M28/14 | 2018.05.31 | 2018 Spring | N47.67119 E20.86590 | SAMN21437879 |
| M28_2018tav_15_2 | M28/15 | 2018.05.31 | 2018 Spring | N47.67134 E20.86593 | SAMN21437880 |
| M28_2018tav_15_3 | M28/15 | 2018.05.31 | 2018 Spring | N47.67134 E20.86593 | SAMN21437881 |
| M28_2018tav_16_2 | M28/16 | 2018.05.31 | 2018 Spring | N47.67101 E20.86648 | SAMN21437882 |
| M28_2018tav_16_3 | M28/16 | 2018.05.31 | 2018 Spring | N47.67101 E20.86648 | SAMN21437883 |
| M28_2018tav_17_1 | M28/17 | 2018.05.31 | 2018 Spring | N47.67098 E20.86695 | SAMN21437884 |
| M28_2018tav_17_2 | M28/17 | 2018.05.31 | 2018 Spring | N47.67098 E20.86695 | SAMN21437885 |
| M28_2018tav_18_2 | M28/18 | 2018.05.31 | 2018 Spring | N47.67083 E20.86577 | SAMN21437886 |
| M28_2018tav_18_3 | M28/18 | 2018.05.31 | 2018 Spring | N47.67083 E20.86577 | SAMN21437887 |
| M28_2018tav_19_2 | M28/19 | 2018.05.31 | 2018 Spring | N47.67099 E20.86771 | SAMN21437888 |
| M28_2019osz_1_1 | M28/1 | 2019.09.24 | 2019 Autumn | N47.67070 E20.86588 | SAMN21437889 |
| M28_2019osz_1_2 | M28/1 | 2019.09.24 | 2019 Autumn | N47.67070 E20.86588 | SAMN21437890 |
| M28_2019osz_1_4 | M28/1 | 2019.09.24 | 2019 Autumn | N47.67070 E20.86588 | SAMN21437891 |
| M28_2019osz_1_5 | M28/1 | 2019.09.24 | 2019 Autumn | N47.67070 E20.86588 | SAMN21437892 |
| M28_2019osz_4_2 | M28/4 | 2019.09.24 | 2019 Autumn | N47.67110 E20.86570 | SAMN21437893 |
| M28_2019osz_4_5 | M28/4 | 2019.09.24 | 2019 Autumn | N47.67110 E20.86570 | SAMN21437894 |
| M28_2019osz_5_1 | M28/5 | 2019.09.24 | 2019 Autumn | N47.67123 E20.86589 | SAMN21437895 |
| M28_2019osz_5_2 | M28/5 | 2019.09.24 | 2019 Autumn | N47.67123 E20.86589 | SAMN21437896 |
| M28_2019osz_5_4 | M28/5 | 2019.09.24 | 2019 Autumn | N47.67123 E20.86589 | SAMN21437897 |
| M28_2019osz_6_1 | M28/6 | 2019.09.24 | 2019 Autumn | N47.67138 E20.86588 | SAMN21437898 |
| M28_2019osz_6_2 | M28/6 | 2019.09.24 | 2019 Autumn | N47.67138 E20.86588 | SAMN21437899 |
| M28_2019osz_6_3 | M28/6 | 2019.09.24 | 2019 Autumn | N47.67138 E20.86588 | SAMN21437900 |
| M28_2019osz_6_5 | M28/6 | 2019.09.24 | 2019 Autumn | N47.67138 E20.86588 | SAMN21437901 |
| M28_2019osz_7_1 | M28/7 | 2019.09.24 | 2019 Autumn | N47.67149 E20.86570 | SAMN21437902 |
| M28_2019osz_7_3 | M28/7 | 2019.09.24 | 2019 Autumn | N47.67149 E20.86570 | SAMN21437903 |
| M28_2019osz_7_4 | M28/7 | 2019.09.24 | 2019 Autumn | N47.67149 E20.86570 | SAMN21437904 |
| M28_2019osz_8_3 | M28/8 | 2019.09.24 | 2019 Autumn | N47.67082 E20.86580 | SAMN21437905 |
| M28_2019osz_8_4 | M28/8 | 2019.09.24 | 2019 Autumn | N47.67082 E20.86580 | SAMN21437906 |
| M28_2019osz_9_1 | M28/9 | 2019.09.24 | 2019 Autumn | N47.67103 E20.86584 | SAMN21437907 |
| M28_2019osz_9_4 | M28/9 | 2019.09.24 | 2019 Autumn | N47.67103 E20.86584 | SAMN21437908 |
| M28_2019osz_10_1 | M28/10 | 2019.09.24 | 2019 Autumn | N47.67121 E20.86594 | SAMN21437909 |
| M28_2019osz_10_2 | M28/10 | 2019.09.24 | 2019 Autumn | N47.67121 E20.86594 | SAMN21437910 |
| M28_2019osz_10_3 | M28/10 | 2019.09.24 | 2019 Autumn | N47.67121 E20.86594 | SAMN21437911 |
| M28_2019osz_10_4 | M28/10 | 2019.09.24 | 2019 Autumn | N47.67121 E20.86594 | SAMN21437912 |
| M28_2019osz_11_2 | M28/11 | 2019.09.24 | 2019 Autumn | N47.67138 E20.86588 | SAMN21437913 |
| M28_2019osz_11_3 | M28/11 | 2019.09.24 | 2019 Autumn | N47.67138 E20.86588 | SAMN21437914 |
| M28_2019osz_12_1 | M28/12 | 2019.09.24 | 2019 Autumn | N47.67113 E20.86642 | SAMN21437915 |
| M28_2019osz_12_2 | M28/12 | 2019.09.24 | 2019 Autumn | N47.67113 E20.86642 | SAMN21437916 |
| M28_2019osz_13_1 | M28/13 | 2019.09.24 | 2019 Autumn | N47.67107 E20.86780 | SAMN21437917 |
| M28_2019osz_13_3 | M28/13 | 2019.09.24 | 2019 Autumn | N47.67107 E20.86780 | SAMN21437918 |
| M28_2019tav_1_3 | M28/1 | 2019.05.16 | 2019 Spring | N47.67120 E20.86294 | SAMN21437919 |
| M28_2019tav_2_1 | M28/2 | 2019.05.16 | 2019 Spring | N47.67119 E20.86292 | SAMN21437920 |
| M28_2019tav_3_1 | M28/3 | 2019.05.16 | 2019 Spring | N47.67120 E20.86305 | SAMN21437921 |
| M28_2019tav_3_2 | M28/3 | 2019.05.16 | 2019 Spring | N47.67120 E20.86305 | SAMN21437922 |
| M28_2019tav_4_1 | M28/4 | 2019.05.16 | 2019 Spring | N47.67115 E20.86312 | SAMN21437923 |
| M28_2019tav_5_1 | M28/5 | 2019.05.16 | 2019 Spring | N47.67110 E20.86330 | SAMN21437924 |
| M28_2019tav_5_2 | M28/5 | 2019.05.16 | 2019 Spring | N47.67110 E20.86330 | SAMN21437925 |
| M28_2019tav_5_3 | M28/5 | 2019.05.16 | 2019 Spring | N47.67110 E20.86330 | SAMN21437926 |
| M28_2019tav_6_2 | M28/6 | 2019.05.16 | 2019 Spring | N47.67105 E20.86331 | SAMN21437927 |
| M28_2019tav_6_3 | M28/6 | 2019.05.16 | 2019 Spring | N47.67105 E20.86331 | SAMN21437928 |
| M28_2019tav_7_1 | M28/7 | 2019.05.16 | 2019 Spring | N47.67110 E20.86379 | SAMN21437929 |
| M28_2019tav_8_2 | M28/8 | 2019.05.16 | 2019 Spring | N47.67103 E20.86385 | SAMN21437930 |
| M28_2019tav_9_2 | M28/9 | 2019.05.16 | 2019 Spring | N47.67103 E20.86402 | SAMN21437931 |
| M28_2019tav_9_3 | M28/9 | 2019.05.16 | 2019 Spring | N47.67103 E20.86402 | SAMN21437932 |
| M28_2019tav_10_3 | M28/10 | 2019.05.16 | 2019 Spring | N47.67111 E20.86420 | SAMN21437933 |
| M28_2019tav_11_1 | M28/11 | 2019.05.16 | 2019 Spring | N47.67112 E20.86434 | SAMN21437934 |
| M28_2019tav_11_2 | M28/11 | 2019.05.16 | 2019 Spring | N47.67112 E20.86434 | SAMN21437935 |
| M28_2019tav_13_2 | M28/13 | 2019.05.16 | 2019 Spring | N47.67106 E20.86496 | SAMN21437936 |
| M28_2019tav_13_3 | M28/13 | 2019.05.16 | 2019 Spring | N47.67106 E20.86496 | SAMN21437937 |
| M28_2019tav_14_1 | M28/14 | 2019.05.16 | 2019 Spring | N47.67133 E20.86584 | SAMN21437938 |
| M28_2019tav_14_2 | M28/14 | 2019.05.16 | 2019 Spring | N47.67133 E20.86584 | SAMN21437939 |
| M28_2019tav_14_3 | M28/14 | 2019.05.16 | 2019 Spring | N47.67133 E20.86584 | SAMN21437940 |
| M28_2019tav_15_1 | M28/15 | 2019.05.16 | 2019 Spring | N47.67078 E20.86577 | SAMN21437941 |
| M28_2019tav_15_2 | M28/15 | 2019.05.16 | 2019 Spring | N47.67078 E20.86577 | SAMN21437942 |
| M28_2019tav_15_3 | M28/15 | 2019.05.16 | 2019 Spring | N47.67078 E20.86577 | SAMN21437943 |
| M28_2019tav_16_1 | M28/16 | 2019.05.16 | 2019 Spring | N47.67079 E20.86584 | SAMN21437944 |
| M28_2019tav_16_2 | M28/16 | 2019.05.16 | 2019 Spring | N47.67079 E20.86584 | SAMN21437945 |
| M28_2019tav_16_3 | M28/16 | 2019.05.16 | 2019 Spring | N47.67079 E20.86584 | SAMN21437946 |
| M28_2019tav_17_1 | M28/17 | 2019.05.16 | 2019 Spring | N47.67096 E20.86584 | SAMN21437947 |
| M28_2019tav_17_2 | M28/17 | 2019.05.16 | 2019 Spring | N47.67096 E20.86584 | SAMN21437948 |
| M28_2019tav_17_3 | M28/17 | 2019.05.16 | 2019 Spring | N47.67096 E20.86584 | SAMN21437949 |
| M28_2019tav_18_1 | M28/18 | 2019.05.16 | 2019 Spring | N47.67135 E20.86592 | SAMN21437950 |
| M28_2019tav_18_3 | M28/18 | 2019.05.16 | 2019 Spring | N47.67135 E20.86592 | SAMN21437951 |
| M28_2019tav_19_1 | M28/19 | 2019.05.16 | 2019 Spring | N47.67091 E20.86687 | SAMN21437952 |
| M28_2019tav_19_2 | M28/19 | 2019.05.16 | 2019 Spring | N47.67091 E20.86687 | SAMN21437953 |
| M28_2019tav_19_3 | M28/19 | 2019.05.16 | 2019 Spring | N47.67091 E20.86687 | SAMN21437954 |
| M28_2019tav_20_1 | M28/20 | 2019.05.16 | 2019 Spring | N47.67102 E20.86720 | SAMN21437955 |
| M28_2019tav_20_2 | M28/20 | 2019.05.16 | 2019 Spring | N47.67102 E20.86720 | SAMN21437956 |

Supplementary Figures:

Supplementary Fig. 1. Distribution of GC content across 1.6 million RAD loci found by the Stacks *de novo* pipeline using default parameters (a). The histogram shows a secondary GC peak likely representing bacterial contamination. RAD loci were aligned to the NCBI *nt* database with *blastn* with an E-value cutoff set to 1e-05. Any locus with a hit to non-cnidarian sequences was considered a contaminant and filtered out to form a contaminants database. Without these contaminants the secondary GC peak was substantially lower (b). The top hits were to *Anthoathecata* (the cnidarian order to which *Hydra* belongs), three bacterial orders (*Burkholderiales*, *Pseudomonadals* and *Aeromonadales*) and *Anostraca* (most likely originating from hydra food).

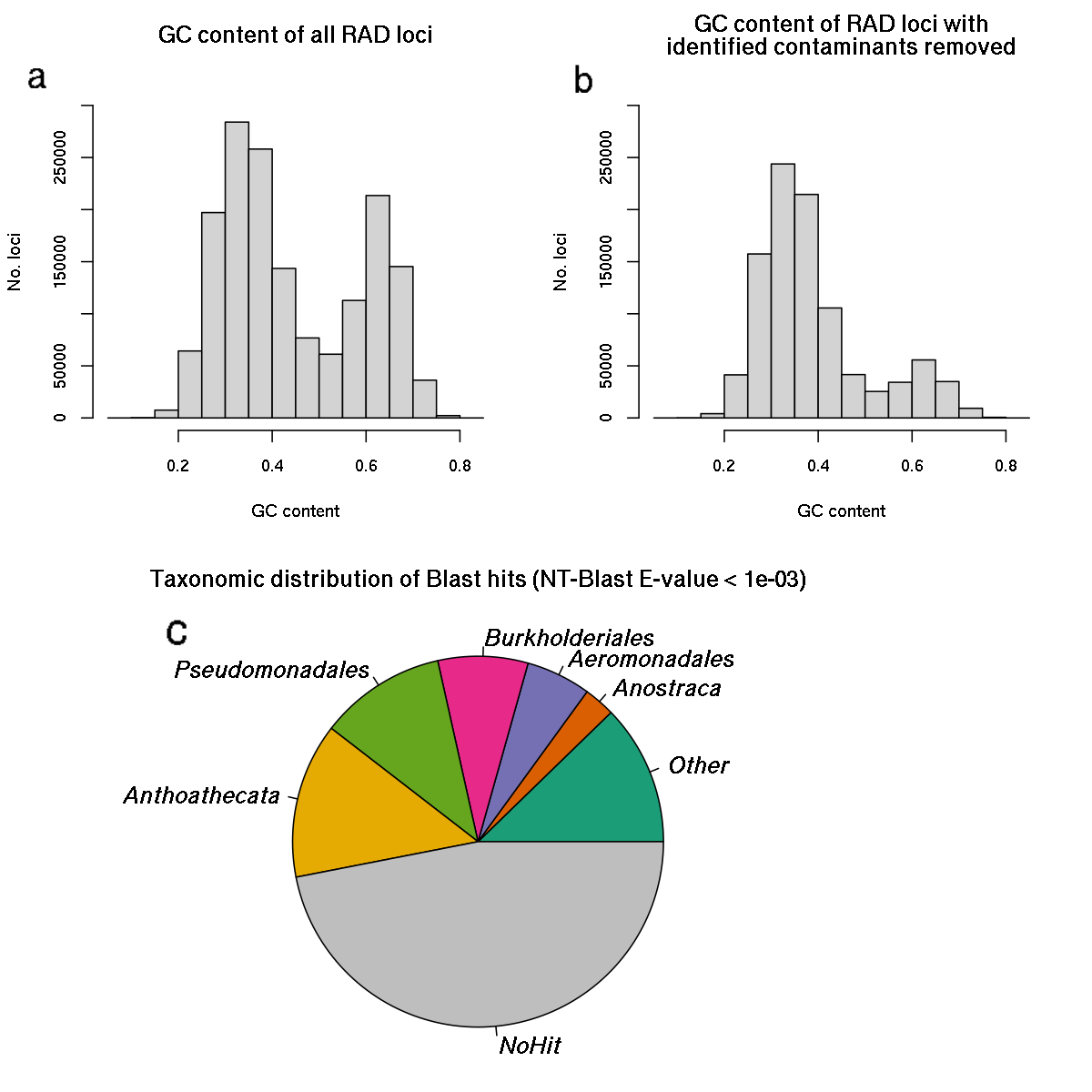

Supplementary Fig. 2. Spectrum of genetic diversity of N=132 H. oligactis strains showing a clear peak of low genetic relatedness (<~0.06, supposed clones) and a secondary peak ~0.11, which can belong to other multilineage genotypes assuming a high genotyping error rate / somatic mutation rate, or distinct multilineage genotypes.

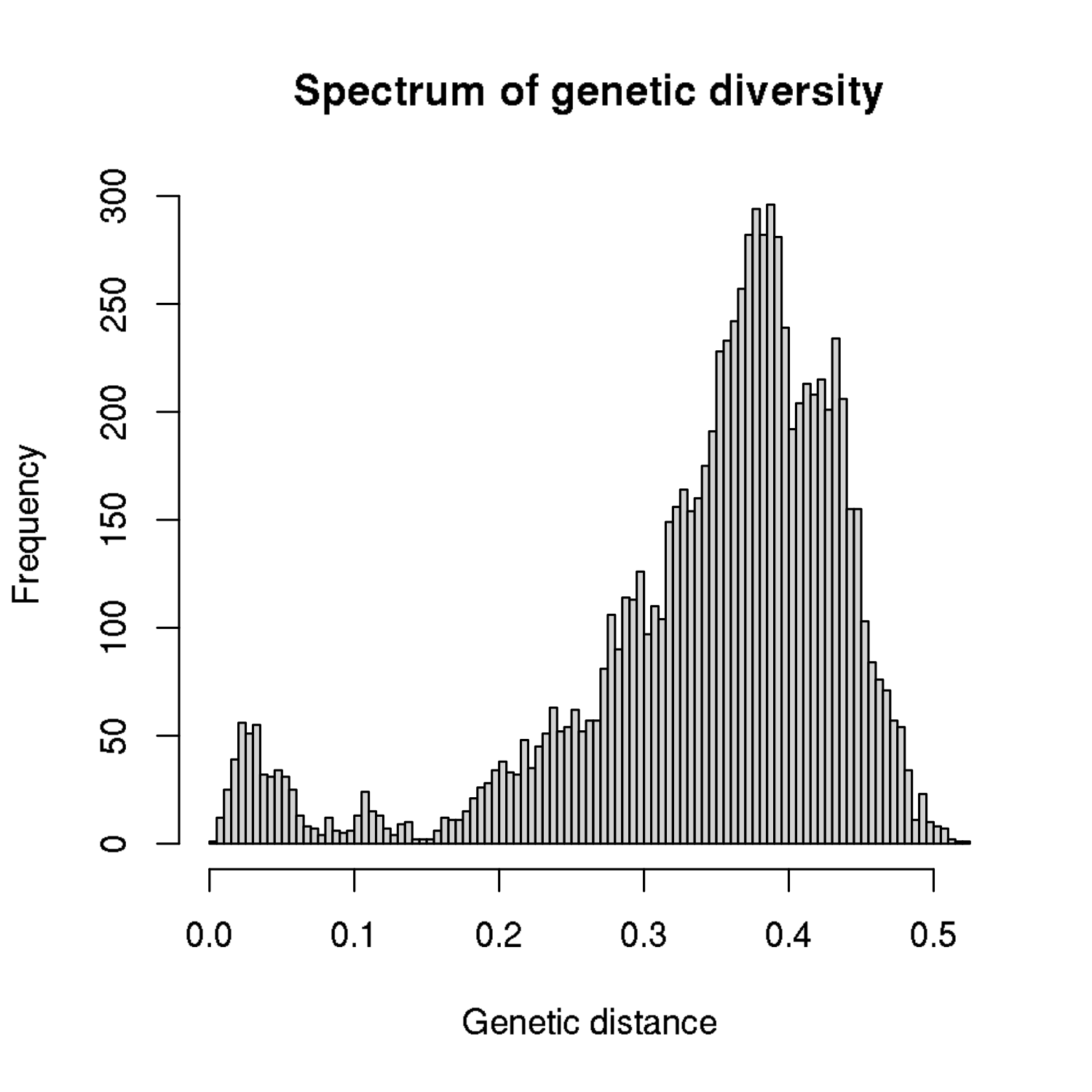
